# Tumor-stromal IL-1β/ NFκB / ESE3 Signaling Axis Drives Pancreatic Cancer Fibrosis, Chemoresistance, and Poor Prognosis

**DOI:** 10.1101/2020.07.27.222455

**Authors:** Tiansuo Zhao, Di Xiao, Fanjie Jin, Hongwei Wang, Jing Liu, Wenrun Cai, Chongbiao Huang, Xiuchao Wang, Song Gao, Shengyu Yang, Jihui Hao

## Abstract

Pancreatic stellate cells (PSCs) play a pivotal role in pancreatic fibrosis and pancreatic ductal adenocarcinoma (PDAC) progression. The mechanisms controlling PSC activation is not completely understood. Here we investigated the role of ESE3 (Epithelium-Specific ETS factor 3) in PSC activation. We discovered that in PDAC patients ESE3 expression was increased in PSC while decreased in tumor cells. ESE3 overexpression in PSC promoted PSC activation. Condition medium from ESE3-overexprssing PSC promotes PDAC cell migration, chemoresistance, tumor growth and fibrosis. ESE3 directly induced the transcription of α-SMA, Collagen 1 and IL-1β by binding to ESE3 binding sites on their promoters to activate PSC. On the other hand, IL-1β upregulates ESE3 in PSC through NFκB activation and ESE3 is required for PSC activation by tumor cell derived IL-1β. Clinical data showed ESE3 upregulation in PSC was positively correlated with tumor size, pTNM stage, CA19-9, CEA and CA242 level in serum. ESE3 overexpression in PSC was an independent negative prognostic factor for disease-free survival and overall survival among PDAC patients. Inhibition of the IL-1β/ ESE3 (PSC)/ IL-1β positive feedback loop represents a promising therapeutic strategy to reduce tumor fibrosis and increase chemotherapeutic efficacy in PDAC.

## Introduction

Pancreatic ductal adenocarcinoma (PDAC) is a highly lethal tumor. Despite recent advances in combination chemotherapy regimens, the 5-year survival rate remains no more than 9% ^(Siegel *et al*, 2020)^. Understanding the molecular mechanisms underlying pancreatic cancer progression and identification of new therapeutic target are urgently needed to improve patient outcome. ESE3 (epithelium-specific ETS factor family member 3) is a member of the ETS superfamily ^(Kas *et al*, 2000)^. ESE3 is highly expressed in normal human pancreas and prostate epithelium ^(Kas *et al*., 2000; Sementchenko & Watson, 2000)^. In prostate cancer, reduced expression of ESE3 was associated with increased expression of epithelial-to-mesenchymal transition (EMT) and stem cell-like genes and clinical features of aggressive disease ^(Albino *et al*, 2016; Albino *et al*, 2012; Dallavalle *et al*, 2016)^. In pancreatic cancer, our previous investigation identified ESE3 as a tumor suppressor. ESE3 overexpression in tumor cells transcriptionally up-regulates E-cadherin to prevent EMT and metastasis ^(Zhao *et al*, 2017)^. Moreover, ESE3 overexpression in PDAC cells down-regulates TGFβ1 and GM-CSF ^(Liu *et al*, 2019)^ to inhibit Treg induction and MDSC accumulation, which in turn increase sensitivity to immunotherapy.

Previous reports support that ESE3 expression in epithelial and tumor cells suppress tumor progression. The functions of ESE3 in stromal cells are not clear. It is reported that ESE3 also expressed in mast cells and monocyte ^(Appel *et al*, 2006; Yamazaki *et al*, 2015)^. In pancreatic cancer only 10-40% of the tumor is consisted of pancreatic cancer cells. The remaining part of the tumor is predominantly desmoplastic stromal cells consisting of pancreatic stellate cells (PSCs) ^(Wilson *et al*, 2014)^. Activated PSCs are the main source of extracellular matrix (ECM) accumulation and fibrosis in pancreatic cancer. PSC plays an important role in promoting PDAC malignancy, chemoresistance and remodeling the tumor microenvironment ^(Amrutkar *et al*, 2019; Cannon *et al*, 2018; Incio *et al*, 2016; Ireland *et al*, 2016)^. It is reported that chronic inflammation plays a crucial role in PSC activation and PDAC fibrosis ^(Bynigeri *et al*, 2017)^. IL-1β, a member of the IL1 family of pro-inflammatory cytokines, has been implicated in PSC activation and pancreatic cancer progression ^(Das *et al*, 2020b; Mitsunaga *et al*, 2013; Zhang *et al*, 2018)^. The mechanisms underlying IL-1β-induced PSC activation is not completely elucidated.

In the present study, we discovered that ESE3 expression in PSC inversely correlated with its expression in pancreatic ductal epithelial cell and PDAC cells. ESE3 overexpression in PSC is crucial for IL-1β induced PSC activation and PDAC fibrosis. Mechanistically, ESE3 activates PSC by directly up-regulating the transcription of α-SMA, collagen-I and IL-1β. On the other hand, IL-1β was able to promote ESE3 overexpression in PSC through NF-κB activation. Importantly, stromal expression of ESE3 is an independent predictor of PDAC progression and survival. Our findings revealed a novel IL-1β /ESE3 / IL-1β feedback signaling loop in PSC activation, PDAC fibrosis and progression.

## Results

### ESE3 was overexpressed in activated PSC

Our previously data demonstrated tumoral ESE3 inhibits PDAC EMT and metastasis by transcriptionally up-regulating E-cadherin ^(Zhao *et al*., 2017)^. Interestingly, although ESE3 expression was down-regulated in PDAC cells, we observed robust upregulation of ESE3 in PSC in PDAC tissues when compared to normal pancreatic tissue (Fig. 1A). Immunofluorescence (IF) staining of ESE3 in human PDAC tissues indicated that ESE3 protein was expressed not only in PDAC cells but also in PSCs, which was consistent with the IHC findings (Fig. 1B).

**Figure 1.**
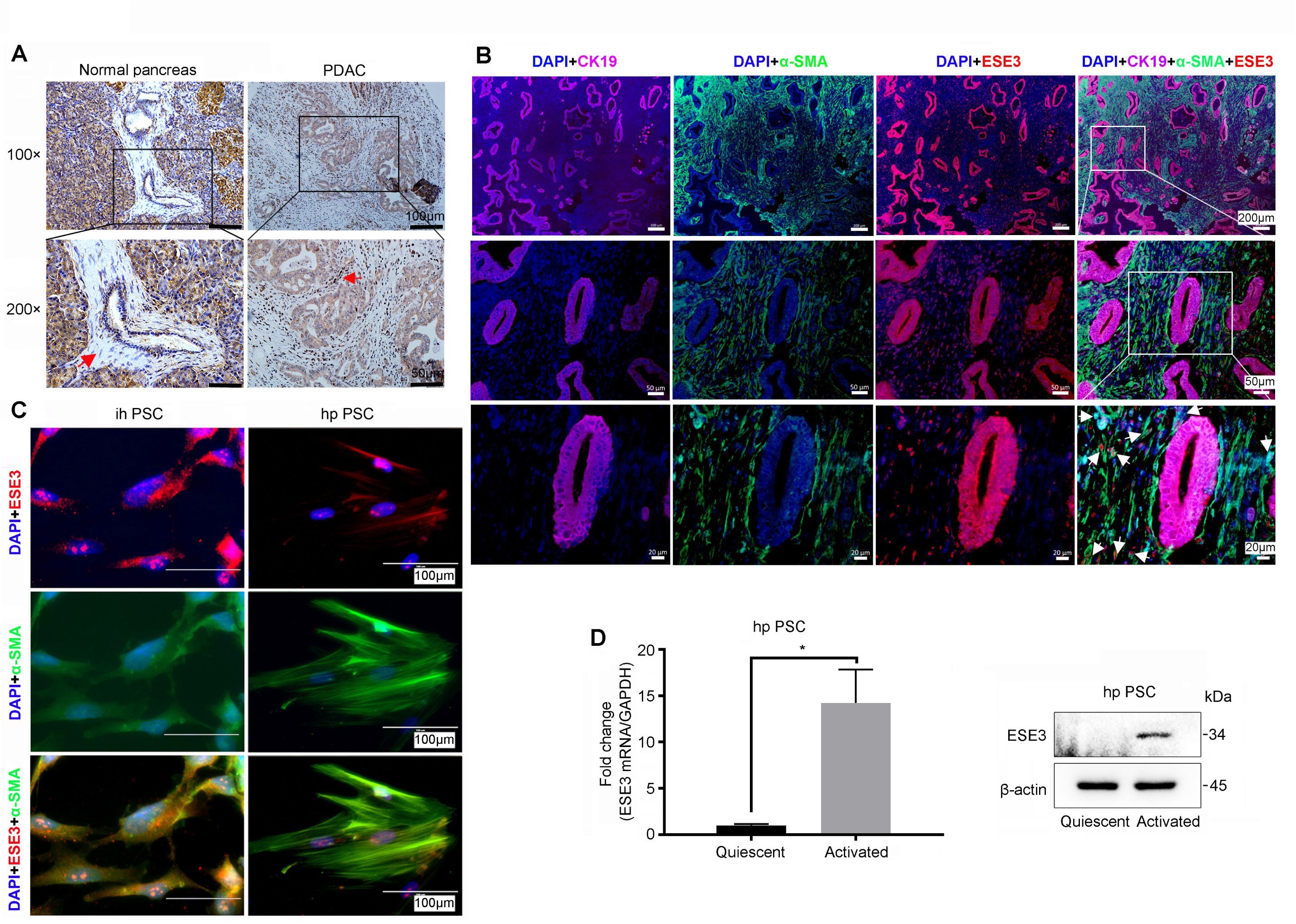
ESE3 was overexpressed in activated PSC. **A**, Immunohistochemical analysis of ESE3 protein expression in PSC of PDAC and adjacent normal pancreatic tissues. **B and C**, IF staining of ESE3, α-SMA (stromal marker) and cytokeratin (CK-19, epithelial marker) expression in PDAC samples (**B**), human primary PSC (hpPSC) and immortal human PSC (ihPSC) (**C**). **D**, the mRNA (left) and protein (right) expression of ESE3 in quiescent and activated hpPSC. \**P* < 0.05. Data information: Data are shown as mean ± SD. P values were determined by two-tailed, two-sample Student’s t-test (D). Experiments were repeated with three biological replicates with similar results.

Furthermore, IF assays were performed in human primary PSC (hpPSC) and immortalized human PSC (ihPSC) to verify ESE3 (PSC) expression and location. ESE3 expression can be found both in cytoplasm and nucleus of PSC (Fig. 1C). To explore the ESE3 expression in quiescent and activated PSC, we performed the PCR and Western-blot assays. The data indicated that ESE3 mRNA and protein expression were significantly elevated in activated PSC compared with its quiescent counterpart (Fig. 1D). Taken together, our data supported that ESE3 was overexpressed in activated PSC in PDAC.

### ESE3 (PSC) overexpression promoted pancreatic cancer progression in vitro and in vivo

To determine whether ESE3 (PSC) plays a role in PDAC progression, ihPSC with ESE3 overexpressed was constructed (pCDH-ESE3 ihPSC). ESE3 expression level was confirmed by the Western-blot (Fig. S1). PDACs were indirectly co-cultured with pCDH-ESE3 ihPSC or pCDH-Vector ihPSC, and the effects of co-culture on PDAC cell apoptosis, cell cycle, cell migration were analyzed. Gem was used to induce PDACs (MIA-PaCa-2 and BxPC-3) apoptosis and G1 phase cell cycle arrest. Both of the apoptosis rate and the G1 phase cell cycle arrest of PDACs treated with Gem was reduced after co-cultured with pCDH-ESE3 ihPSC compared with those co-cultured with pCDH-Vector ihPSC (Fig. 2A and B). Trans-well and wound-healing assays demonstrated that CM collected from pCDH-ESE3 ihPSC significantly promoted the PDAC cell migration (Fig. 2C and D). Conversely, ESE3 knockdown in ihPSC significantly reduced the ability of ihPSC to promote PDAC cell migration in co-culture experiments (Fig. S2). Taken together, our data indicated that ESE3 (PSC) overexpression is crucial for PSC to promote PDACs survival, chemoresistance and cell migration.

**Figure 2.**
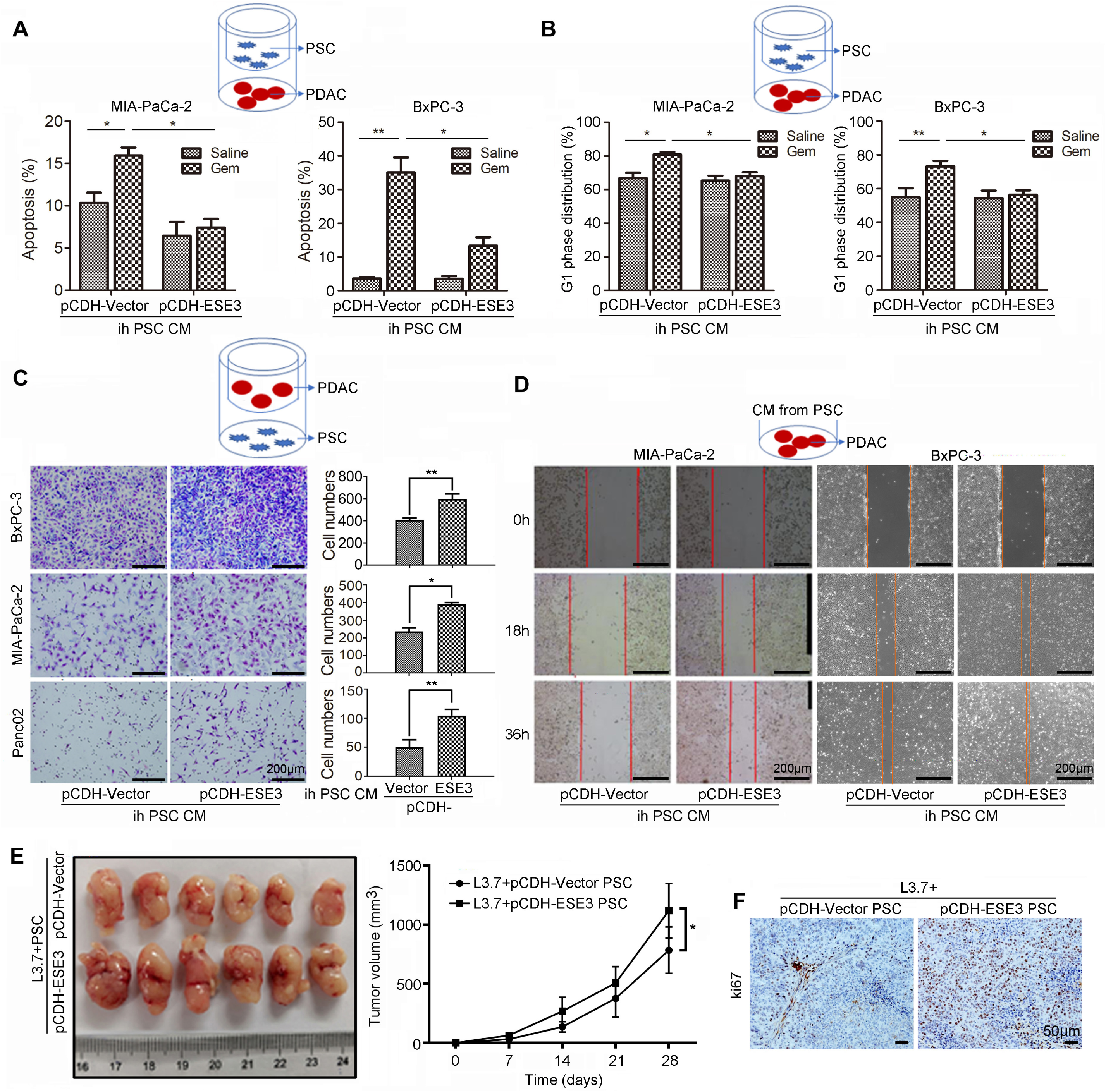
ESE3 (PSC) overexpression promoted pancreatic cancer progression in vitro and in vivo. **A and B**, Flow cytometry to analyze the apoptosis (**A**) and G1 phase distribution (**B**) of MIA-PaCa-2 and BxPC-3 cells indirectly co-cultured with pCDH-ESE3 ihPSC or pCDH-Vector ihPSC and then treated with Gem (2 μM) for 24 hours. **C**, Comparison of the migration of MIA-PaCa-2, BxPC-3 and Pan02 cells indirectly co-cultured with pCDH-ESE3 ihPSC or pCDH-Vector ihPSC for 18 hours by Trans-well assay. **D**, Wound-healing assays comparing the motility of MIA-PaCa-2 and BxPC-3 cells cultured with CM from pCDH-ESE3 ihPSC or pCDH-Vector ihPSC. NC means negative control. **E**, Representative images of mice injected with ihPSC and PDAC cells (1:1) subcutaneously implanted into the nude mice (nu/nu) (left). The data on all primary tumors are expressed as mean ± SD (right). **F**, IHC analysis of ki67 expression in the mouse xenograft samples. \**P* < 0.05 and \*\**P* < 0.01. Data information: Data are shown as mean ± SD. Experiments in (A, B, C and D) were repeated with three biological replicates with similar results. n = 6 in (E). P values in were determined by two-tailed, two-sample Student’s t-test.

To evaluate whether ESE3 (PSC) overexpression drives PDAC progression in vivo, PSC and L3.7 cell mixture (1:1) were subcutaneously injected into both flanks of nude nu/nu mice. Compared with the control group (pCDH-Vector ihPSC and L3.7 cell), the average tumor volume in the experiment group (pCDH-ESE3 ihPSC and L3.7 cell) was significantly increased (Fig. 2E). IHC was used to verify the expression of ki67 in tumor and PSC using the mouse xenograft samples. ki67 in tumor and PSC were both elevated significantly in the experiment group compared with the control group (Fig. 2F).

### ESE3 induced α-SMA, Collagen1 and IL-1β expression in PSC

Elevated expression α-SMA, accumulation of extracellular matrix (ECM) protein (such as collagens) and increased secretion inflammatory cytokines in PSC are markers of pancreatic fibrosis in chronic pancreatitis and pancreatic cancer ^(Xue *et al*, 2018)^. To investigate the role of ESE3 overexpression in PSC activation, we examine the effect of ectopic ESE3 on the expression of α-SMA, Collagen1 and IL-1β. pCDH-ESE3 plasmid was transfected in ihPSC, human primary PSC (hpPSC) and mouse primary PSC (mpPSC). PCR and western-blot demonstrated that the mRNA and protein level of α-SMA, Collagen1 and IL-1β were significantly upregulated by ectopically expressed ESE3 (Fig. 3A-C).

**Figure 3.**
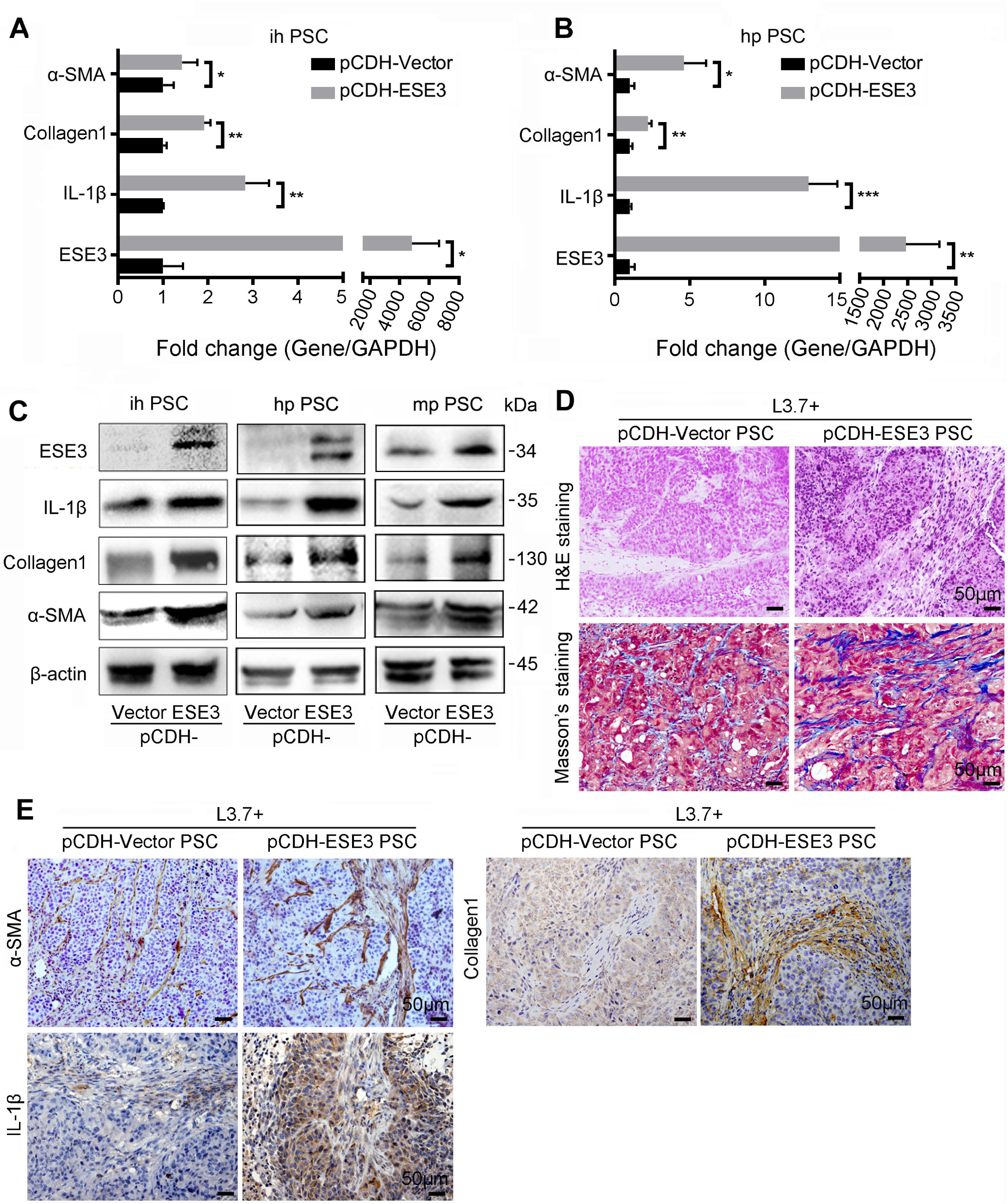
ESE3 induced α-SMA, Collagen1 and IL-1β expression in PSC. **A and B**, mRNA expression of α-SMA, Collagen1 and IL-1β in ihPSC (**A**) and hpPSC (**B**) transfected with pCDH-ESE3 or pCDH-Vector plasmids. **C**, Protein expression of α-SMA, Collagen1 and IL-1β in ihPSC, hpPSC and mpPSC transfected with pCDH-ESE3 or pCDH-Vector plasmids. **D**, HȆE staining and masson’s trichrome staining in the mouse xenograft samples. **E**, IHC analysis of α-SMA, IL-1β and Collagen 1 expression in the mouse xenograft samples. \**P* < 0.05, \*\**P* < 0.01 and \*\*\**P* < 0.001. Data information: Data are shown as mean ± SD. Experiments in (A and B) were repeated with three biological replicates with similar results. P values in were determined by two-tailed, two-sample Student’s t-test.

For in vivo study, PSC and L3.7 cell (1:1) were subcutaneously injected into the two flanks of nude nu/nu mice. The mouse xenograft samples were furtherly used to evaluate the PSC activation status. H&E and masson’s trichrome staining indicated that the experiment group (pCDH-ESE3 ihPSC and L3.7 cell) has more fibrosis than the control group (pCDH-Vector ihPSC and L3.7 cell) (Fig. 3D). The expression of α-SMA, IL-1β and Collagen1 in PSC were also significantly increased in the experiment group (Fig. 3E). Taken together, these results suggested that ESE3 induced PSC activation in vitro and aggravated the fibrotic reaction in xenograft tumors.

### ESE3 directly binds to the promoter regions of α-SMA, Collagen1 and IL-1β to upregulate their expression

To understand the molecular mechanism by which ESE3 activates PSC, we surveyed the promoter region of α-SMA, Collagen1 and IL-1β genes and identified ETS-binding sites (EBS; GGAA/T) in the promoters (Fig. 4A, C and E, top). ChIP and luciferase analysis were performed. In chromatin fractions pulled down by an anti-ESE3 antibody, the fragment immunoprecipitated by anti-ESE3 antibody were detected (Fig. 4A, C and E, bottom). To determine whether the binding of ESE3 activates α-SMA, Collagen1 and IL-1β promoter, different α-SMA, Collagen1 and IL-1β luciferase promoter reporter constructs were constructed and co-transfected with pCDH-Vector or pCDH-ESE3 plasmid into HEK 293 cell and ihPSC. Luciferase analysis showed that ESE3 overexpressed by pCDH-ESE3 plasmid significantly increased α-SMA, Collagen1 and IL-1β promoter activity in HEK 293 cell and ihPSC (Fig. 4B, D and F). To determine whether the EBSs are required for ESE3 to transactivate α-SMA, Collagen1 and IL-1β promoter, these EBSs were mutated. As shown in Fig. 4B, D and F, the mutation of EBSs almost abolished the transactivation of α-SMA, Collagen1 and IL-1β promoter by ESE3. These data indicated that ESE3 directly binds to promoters of α-SMA, Collagen1 and IL-1β to induce the expression of these genes and to activate PSC.

**Figure 4.**
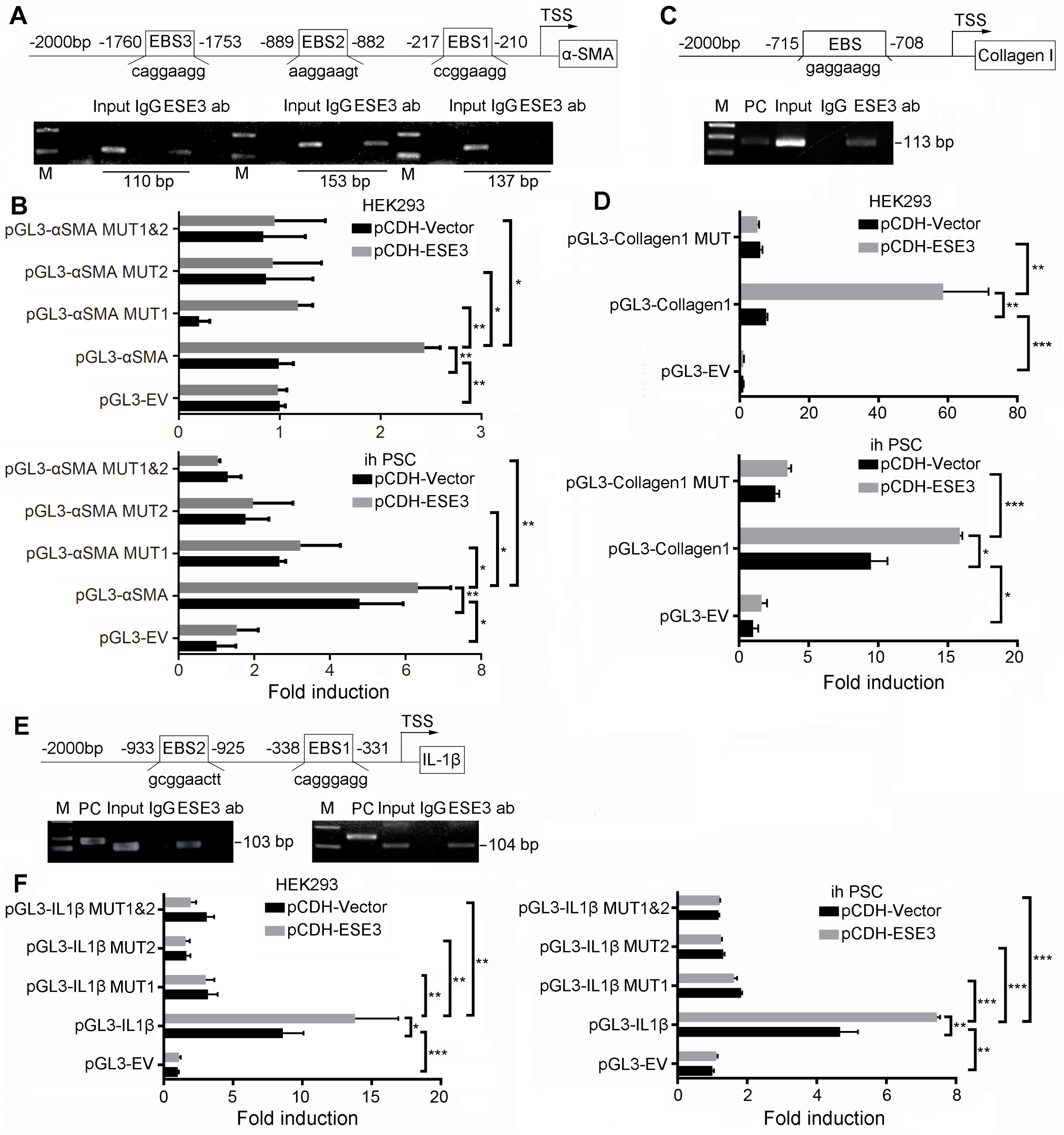
ESE3 directly binds to the promoter regions of α-SMA, Collagen1 and IL-1β to upregulate their expression. **A, C and E**, Schematic of the structure of the α-SMA (**A**), Collagen1 (**C**) and IL-1β (**E**) gene promoters. Shown are EBSs and their location (upper). Chromatin immunoprecipitation analysis of ESE3 binding to the α-SMA (**A**), Collagen1 (**C**) and IL-1β (**E**) promoter in ihPSC (lower). **B, D and F**, Luciferase assay-based promoter activity analysis of HEK293 and ihPSC overexpressed ESE3 (pCDH-ESE3) and control (pCDH-Vector) cells transfected with pGL3-α-SMA (**B**), pGL3-Collagen1 (**D**) and pGL3-IL-1β (**F**), pGL3-Empty Vector (pGL3-EV) and pGL3-Mutations (pGL3-MUT). Forty-eight hours after transfection, the cells were subjected to dual luciferase analysis. The results are expressed as fold induction relative to that in corresponding cells transfected with the control vector after normalization of firefly luciferase activity according to *Renilla* luciferase activity. The data are expressed as the means ± SD from three independent experiments. M means marker. \**P* < 0.05, \*\**P* < 0.01 and \*\*\**P* < 0.001. Data information: Data are shown as mean ± SD. Experiments in (B, D and F) were repeated with three biological replicates with similar results. P values in were determined by two-tailed, two-sample Student’s t-test.

### Tumor secreted IL-1β induced nuclear ESE3 (PSC) expression by activating NF-κB

IL-1β plays an important role in pancreatic fibrosis and PSC activation ^(Das *et al*, 2020a)^. Previous studies demonstrated IL-1β was mainly produced by pancreatic cancer stroma ^(Daley *et al*, 2017; Incio *et al*., 2016; Ochi *et al*, 2012)^. Recent research found that tumor cell-derived IL-1β is essential for the establishment of pancreatic cancer desmoplasia ^(Das *et al*., 2020b)^. We furtherly tested the IL-1β expression in PDAC cell lines treated with or without Gem. Western-blot analysis indicated that tumor derived IL-1β protein increased after Gem treatment (Fig. 5A). ELISA assay demonstrated a significantly increased of secreted IL-1β concentration in the conditional medium (CM) of PDAC cell lines treated with Gem (Fig. 5B). Given the role of IL-1β in PSC activation, we examined the role of IL-1β in ESE3 (PSC) expression, PSC was treated with recombined IL-1β or DMSO. IL-1β treatment was sufficient to induce ESE3 expression in PSCs (Fig. 5C).

**Figure 5.**
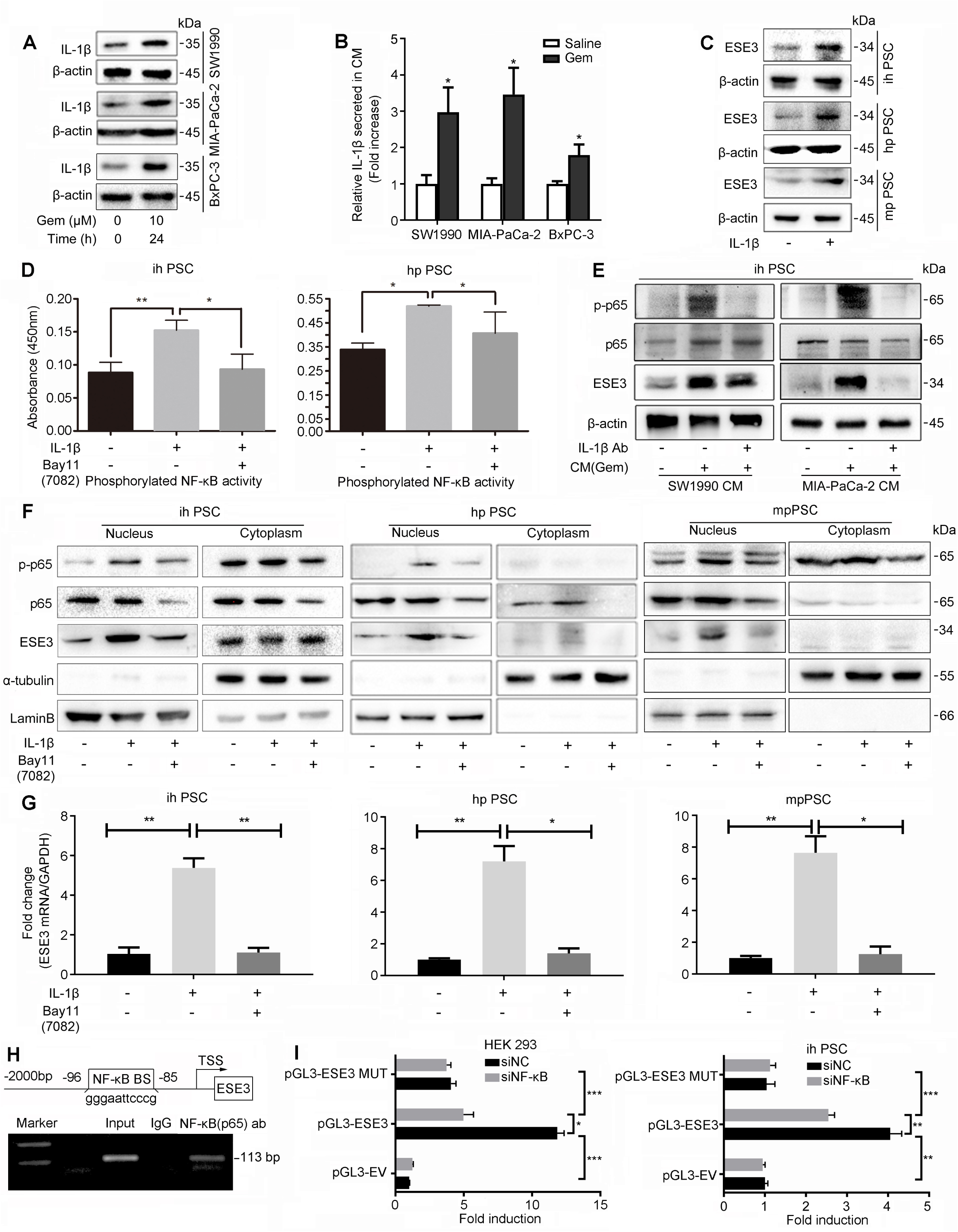
Tumor secreted IL-1β induced nuclear ESE3 (PSC) expression by activating NF-κB. **A**, Western blot analysis of IL-1β expression in SW1990, MIA-PaCa-2 and BxPC-3 cell lines treated with Gem (10 μM) for 24 hours. **B**, ELISA analysis of IL-1β secretion in CM of SW1990, MIA-PaCa-2 and BxPC-3 cell lines treated with Gem (10 μM) for 24 hours. **C**, Western blot analysis of ESE3 in ihPSC, hpPSC and mpPSC treated with recombinant human IL-1β (100 ng/mL, 24 hours). **D**, Analysis of the NF-κB (p65) transcriptional activity in ihPSC and hpPSC treated with recombinant human IL-1β (100ng/mL, 24 hours) or / and NF-κB (p65) inhibitor (Bay11-7082) (8 μM,12 hours) by commercial kit. **E**, Western blot analysis of p-p65 and ESE3 in ihPSC cultured with CM (SW1990 and MIA-PaCa-2 cell lines treated with Gem) or / and IL-1β neutralizing antibody for 24 hours. **F**, Western blot analysis of p-p65 and ESE3 expression in nucleus and cytoplasm separation form the whole protein of PSCs treated with recombinant human IL-1β or / and Bay11-7082. **G**, RT-PCR analysis of ESE3 mRNA expression in ihPSC, hpPSC and mpPSC treated with recombinant human IL-1β or / and Bay11-7082. **H**, Schematic of the structure of the ESE3 gene promoter. Shown is one κB binding site and the location (upper). Chromatin immunoprecipitation analysis of NF-κB binding to the ESE3 promoter in ihPSC (lower). **I**, Luciferase assay-based promoter activity analysis of HEK293 (left) and ihPSC (right) knockdown NF-κB (p65) (si p65) and control cells (si NC) transfected with pGL3-ESE3, pGL3-Empty Vector (pGL3-EV) and pGL3-Mutation (pGL3-MUT). Forty-eight hours after transfection, the cells were subjected to dual luciferase analysis. The results are expressed as fold induction relative to that in corresponding cells transfected with the control vector after normalization of firefly luciferase activity according to Renilla luciferase activity. The data are expressed as the means ± SD from three independent experiments. \**P* < 0.05, \*\**P* < 0.01 and \*\*\**P* < 0.001. Data information: Data are shown as mean ± SD. Experiments in (B, D, G and I) were repeated with three biological replicates with similar results. P values in were determined by two-tailed, two-sample Student’s t-test.

To understand the mechanism by which IL-1 β regulates ESE3, genome-wide mRNA sequencing was performed to find the pathway activated by IL-1β stimulation. Pathway enrichment analysis showed that NF-κB signaling was significantly activated in IL-1β treated PSC (Fig. S3). To determine whether IL-1β induced ESE3 (PSC) expression through activating NF-κB, Bay11-7082 was used to block the NF-κB signal pathway. NF-κB (p65) transcriptional activity in ihPSC and hpPSC treated with or without recombinant human IL-1β was furtherly confirmed using p65 binding ELISA assays. IL-1β significantly activated NF-κB (p65) transcriptional activity, which can be blocked by Bay11-7082 (Fig. 5D).

To determine if tumor derived CM induces ESE3 (PSC) overexpression, ihPSC was treated with CM or non Gem treated CM. The CM is collected from SW1990 and MIA-PaCa-2 cell lines that treated with Gem. Western-blot indicated that the expression of ESE3 was upregulated in ihPSC treated with CM compared with those treated with non Gem treated CM (Fig. 5E). And the CM induced ESE3 (PSC) overexpression can be blocked by IL-1β neutralizing antibody (Fig. 5E). Nuclear p-p65 and ESE3 protein expression was also significantly upregulated in ihPSC, hpPSC and mpPSC treated with recombinant IL-1β (Fig. 5F). IL-1β induced ESE3 overexpression and p65 phosphorylation at protein and mRNA levels were inhibited by p65 inhibitor Bay11-7082, suggesting that IL-1β induced ESE3 expression in PSC through NF-κB (Fig. 5F and G).

### NF-κB regulated ESE3 expression by directly binding to its promoter region

NF-κB regulates target gene expression through binding to promoter and enhancer regions containing κB consensus sequences 5′ GGGRNWYYCC 3′ (N, any base; R, purine; W, adenine or thymine; Y, pyrimidine) ^(Zhang *et al*, 2017)^. We identified a κB binding site in the promoter region of human ESE3 gene (Fig. 5H, top). To examine whether NF-κB (p65) directly binds to ESE3 promoter, ChIP assay was performed in ihPSC. In chromatin fractions pulled down by an anti-p65 antibody, the fragment immunoprecipitated by anti-p65 antibody was detected (Fig. 5H, bottom). To determine whether the binding of NF-κB (p65) activates ESE3 promoter, we constructed a full-length ESE3 luciferase promoter vector and co-transfected this reporter construct with or without siNF-κB (p65) into HEK 293 cell and ihPSC. Luciferase analysis showed that suppressed NF-κB activity by siNF-κB (p65) significantly decreased ESE3 promoter activity in HEK 293 cell and ihPSC (Fig. 5I). To determine whether the κB binding site is required for NF-κB to transactivate ESE3 promoter, this κB binding site was mutated. As shown in Fig. 5I, the mutation of κB binding site almost abolished the transactivation of ESE3 promoter by NF-κB.

### IL-1β/ ESE3(PSCs) / IL-1β positive feedback loop promoted PDAC progression

As shown in Fig. 6A and B, the mRNA and protein expression of ESE3, α-SMA, Collagen1 and IL-1β were induced by recombinant IL-1β, which is consistent with the activation of PSC by IL-1β. To further understand the role of ESE3 in IL-1β mediated PDAC progression, blocking experiment were conducted using PSC and PSC-siESE3. After receiving IL-1β stimulation, the expression of α-SMA, Collagen1 and IL-1β in PSC were significantly upregulated. Conversely, in PSC-siESE3 cell lines, compared with the group without IL-1β stimulation, no change was observed in the expression of α-SMA, Collagen1 and IL-1β in the group which received IL-1β stimulation (Fig. 6C). Furthermore, we tested the role of NF-κB (p65) in the IL-1β induced cascade by using Bay11-7082. When NF-κB (p65) activity was inhibited by Bay11-7082, the protein and mRNA expression of ESE3, α-SMA, Collagen1 and IL-1β were reduced in ihPSC and hpPSC (Fig. 6D and E). Moreover, we used the sip65 to suppress NF-κB (p65) activity and got the same result as using Bay11-7082 for blocking (Fig. 6F).

**Figure 6.**
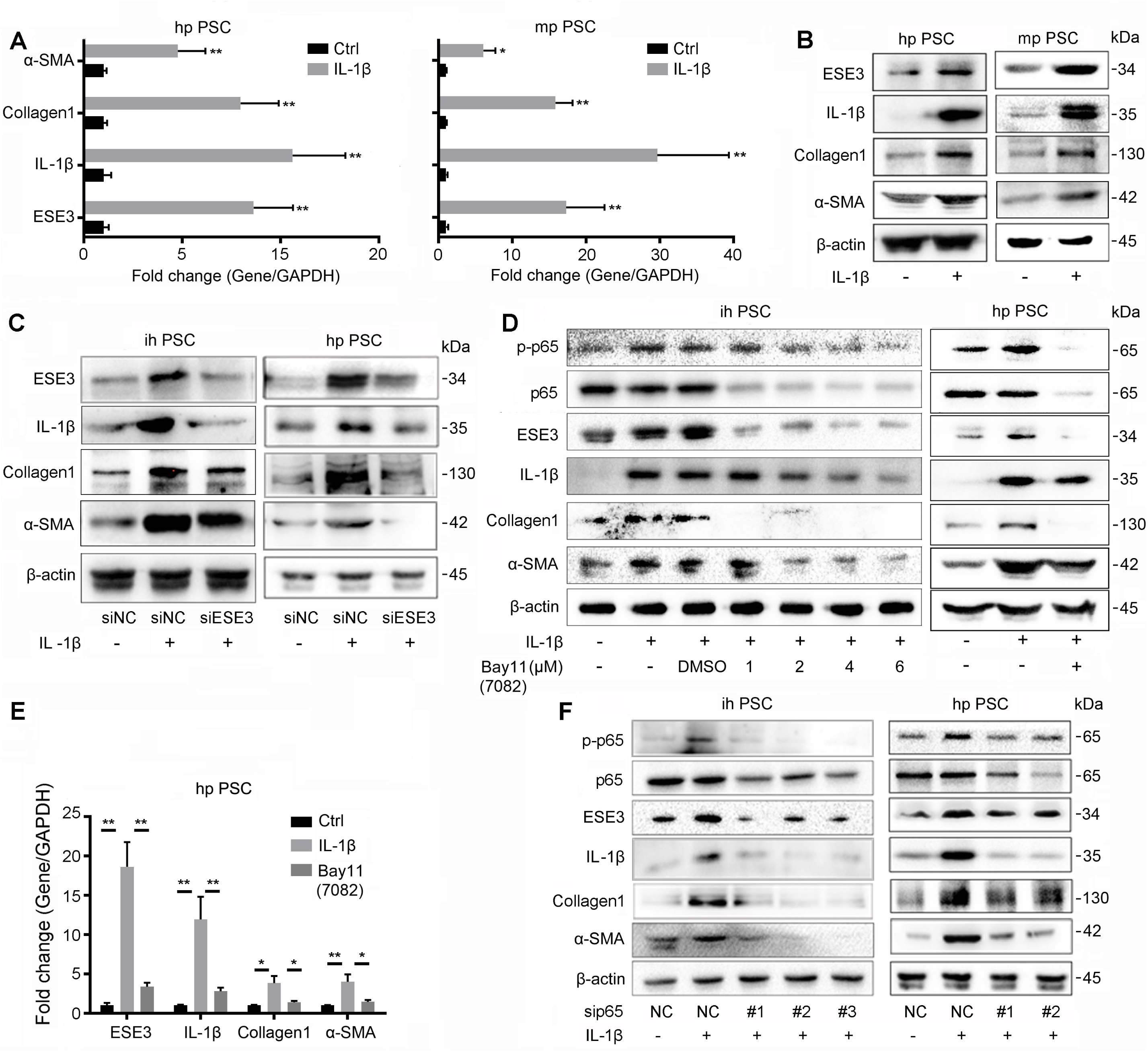
IL-1β/ ESE3(PSCs) / IL-1β positive feedback loop promoted PDAC progression. **A and B**, mRNA (**A**), and protein (**B**) expression of ESE3, α-SMA, Collagen1 and IL-1β in hpPSC and mpPSC stimulated with recombinant human IL-1β (100 ng/mL, 24 hours). **C**, Western blot analysis of ESE3, α-SMA, Collagen1 and IL-1β expression in ihPSC and hpPSC with IL-1β stimulation or/ and knockdown of ESE3 expression. **D and E**, Western blot (**D**) and RT-PCR (**E**) analysis of p-p65, ESE3, α-SMA, Collagen1 and IL-1β expression in ihPSC and hpPSC with IL-1β stimulation or/ and NF-κB (p65) inhibitor Bay11-7082 treatment. **F**, Western blot analysis of p-p65, ESE3, α-SMA, Collagen1 and IL-1β expression in ihPSC and hpPSC with IL-1β stimulation or/ and knockdown of NF-κB (p65) expression. \**P* < 0.05 and \*\**P* < 0.01. Data information: Data are shown as mean ± SD. Experiments in (A and E) were repeated with three biological replicates with similar results. P values in were determined by two-tailed, two-sample Student’s t-test.

### ESE3 (PSC) was associated with decreased DFS and OS in PDAC patients

To understand the significance of stromal ESE3 expression in PDAC progression, ESE3 IHC score was evaluated by two independent pathologists. An inverse correlation can be observed between ESE3 (PSC) expression and ESE3 (tumor) expression (Fig. 7A, *P* = 0. 033, *r* = −0.318). To explore the pathologic significance of ESE3 (PSC) regarding PDAC progression, the correlation between ESE3 (PSC) and clinicopathological features of PDAC was evaluated (Table 1). No significant correlation between ESE3 (PSC) and gender, age, differentiation, lymph node metastasis and histologic grade was observed. However, ESE3 (PSC) was positively correlated with pTNM (𝒳^2^ = 6.343, *P* = 0.012), tumor size (𝒳^2^ = 6.405, *P* = 0.011), CA19-9 (𝒳^2^ = 4.845, *P* = 0.028), CEA (𝒳^2^ = 6.060, *P* = 0.014) and CA242 (𝒳^2^ = 4.046, *P*= 0.044) (Table 1). Importantly, Kaplan-Meier analysis of IHC data indicated that PDAC patients with moderate (++) or high (+++) levels of ES E3 (PSC) had significantly worse DFS and OS than did those with negative (-) or low (+) ESE3 protein expression (*P* < 0.05; DFS: 14.00 and 33.70 months; OS: 17.00 and 33.73 months, respectively) (Fig. 7B). Univariate and multivariate analysis of clinical follow-up data indicated that ESE3 expression in PSC was an independent risk factor for both OS and DFS (Table S1). Taken together, these data demonstrated IL-1β upregulate ESE3 expression in PSC to transactivate α-SMA, Collagen1 and IL-1β to promote PDAC fibrosis. The positive feedback loop of IL-1β/ ESE3(PSC)/ IL-1β promoted PDAC fibrosis, chemoresistance and poor prognosis (Fig. 7C).

**Table 1.** Association between ESE3 expression in PSCs and clinicopathological feature of patients with PDAC tissues

| | Total | ESE3 expression | | $\chi^2$ | P value |
| --- | --- | --- | --- | --- | --- |
|  |  | High/Moderate | Low/absent |  |  |
| Gender |  |  |  | 1.551 | 0.213 |
| Male | 59 | 39 | 20 |  |  |
| Female | 48 | 37 | 11 |  |  |
| Age(year) |  |  |  | 0.587 | 0.444 |
| <65 | 78 | 57 | 21 |  |  |
| ≥65 | 29 | 19 | 10 |  |  |
| Differentiation |  |  |  | 0.001 | 0.981 |
| Well/Moderate | 83 | 59 | 24 |  |  |
| Poor | 24 | 17 | 7 |  |  |
| pTNM |  |  |  | 6.343 | 0.012* |
| I-II | 30 | 16 | 14 |  |  |
| III | 77 | 60 | 17 |  |  |
| Tumor size(cm) |  |  |  | 6.405 | 0.011* |
| <5 | 59 | 36 | 23 |  |  |
| ≥5 | 48 | 40 | 8 |  |  |
| LN metastasis |  |  |  | 0.163 | 0.687 |
| - | 27 | 20 | 7 |  |  |
| + | 80 | 56 | 24 |  |  |
| Histological grade |  |  |  | 0.213 | 0.645 |
| G1 | 76 | 53 | 23 |  |  |
| G2 and G3 | 31 | 23 | 8 |  |  |
| CA19-9 ( kU/L ) |  |  |  |  |  |
| <40 | 32 | 18 | 14 | 4.845 | 0.028* |
| ≥40 | 75 | 58 | 17 |  |  |
| CEA ( μg/L ) |  |  |  |  |  |
| <5 | 67 | 42 | 25 | 6.060 | 0.014* |
| ≥5 | 40 | 34 | 6 |  |  |
| CA242 ( IU/mL ) |  |  |  |  |  |
| <20 | 46 | 28 | 18 | 4.046 | 0.044* |
| ≥20 | 61 | 48 | 13 |  |  |

Abbreviation: LN, lymph node. \* $P < 0.05$ (Chi-Square Test)

**Figure 7.**
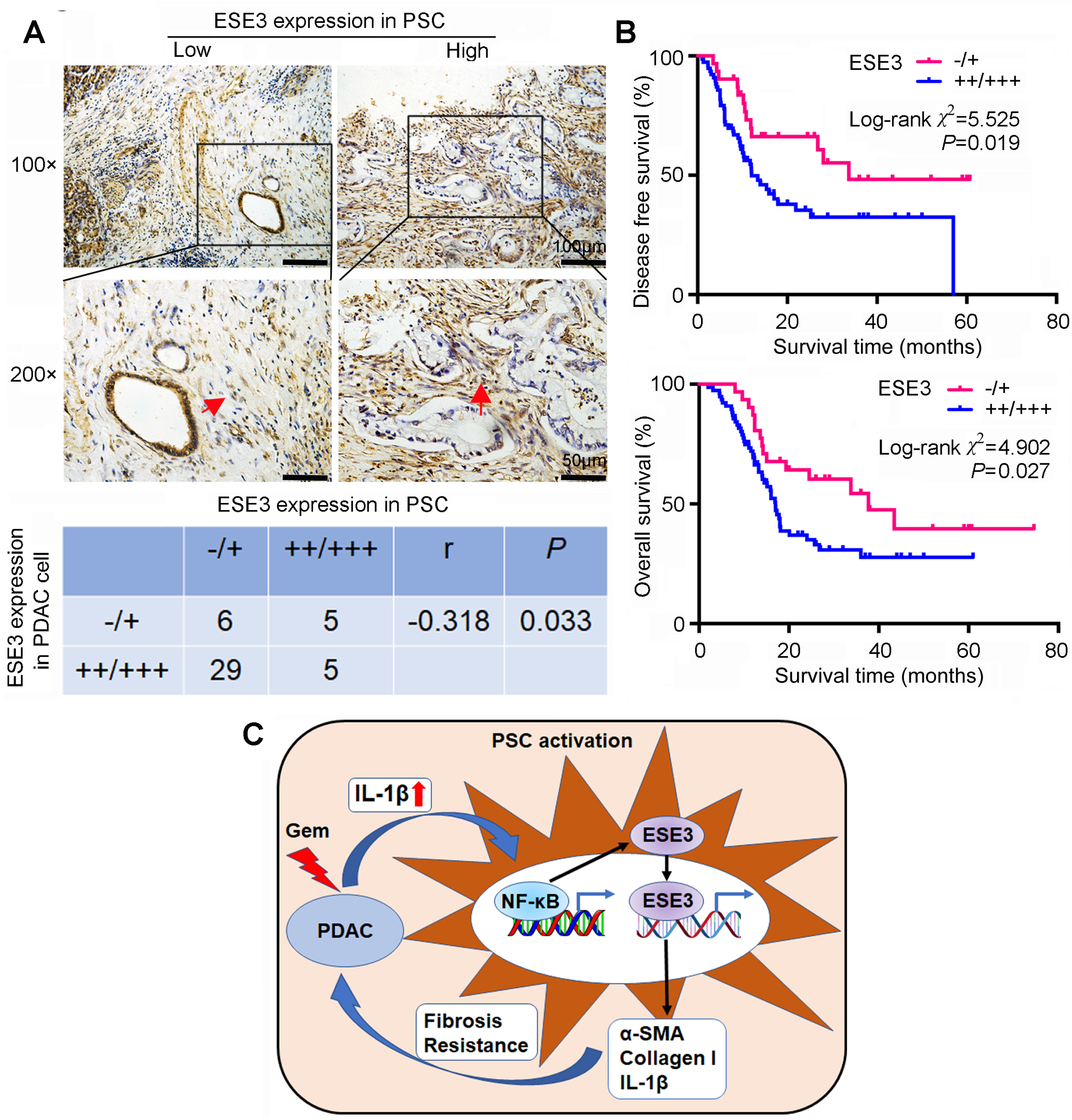
ESE3 (PSC) was associated with decreased DFS and OS in PDAC patients. **A**, Immunohistochemical analysis of ESE3 protein expression in PSCs of PDAC tissues (upper). The relationship of ESE3 expression between PSC and PDAC cells in PDAC samples (lower). r = −0.318, P = 0.033, as determined by rank correlation test. **B**, Association of ESE3 expression in PSC with DFS (upper) and OS (lower) in PDAC patients. PDAC patients with moderate (++) or high (+++) ESE3 expression in PSC had significantly shorter median DFS and OS than those with negative (-) or low (+) ESE3 protein expression (*P* < 0.05). **C**, Schematic of the IL-1β/ NF-κB/ ESE3 signaling axis increased α-SMA, Collagen1 and IL-1β expression in PSC and promoted fibrosis and chemoresistance. Data information: Kaplan-Meier method (log-rank test) was used to estimate the overall survival (OS) or disease-free survival (DFS) of different ESE3 expression in PDAC (B).

## Discussion

Pancreatic cancer is the seventh leading cause of global cancer deaths worldwide. Although the past few years have seen improvements in first-line and second-line palliative therapies, the 5-year survival rate of PDAC remains poor ^(Rawla *et al*, 2019)^.

ESE3 is a member of the highly diverse ETS superfamily, which is a newly identified tumor suppressor. Our group previously reported that tumoral ESE3 is a tumor suppressing transcription factor that directly inhibits PDAC metastasis by up-regulating E-cadherin ^(Zhao *et al*., 2017)^. In addition, tumoral ESE3 inhibited the expression of TGFβ1 and GM-CSF to promote the immune-suppressive tumor microenvironment in PDAC. Tumoral EHF can be used as a promising biomarker to evaluate the immune microenvironment status of PDAC and to screen for patient response to anti-PD1 therapy ^(Liu *et al*., 2019)^.

Previous research suggested that the function of ESE3 is context-dependent ^(Hollenhorst *et al*, 2011; Madison *et al*, 2018)^. Tugores A et al reported ESE3 was a downstream signal of MAPK pathway ^(Tugores *et al*, 2001)^. It is reported that ESE3 is not only play a critical role in tumor cells, but also express in mast cells and monocyte. Furthermore, ESE3 mRNA and protein also express in human bronchial smooth muscle cells ^(Silverman *et al*, 2002)^. In human bronchial smooth muscle cells, ESE-3 mRNA and protein can be upregulated by IL-1β ^(Silverman *et al*., 2002)^. IL-1β plays a key role in carcinogenesis and tumor growth, due to its importance in mediating inflammatory response. Increased levels of IL-1β in body fluids are correlated with worse prognosis, chemoresistance and invasiveness in cancer ^(Zhang *et al*., 2018)^. IL-1β promotes drives chronic inflammation, fibrosis, endothelial cell activation, tumor angiogenesis and induction of immunosuppressive cells to promote tumor progression ^(Incio *et al*., 2016; Mitsunaga *et al*., 2013; Nomura *et al*, 2018; Simpson *et al*, 2019)^. Previous studies posit that IL-1β in the PDAC microenvironment was mainly produced by the stroma ^(Daley *et al*., 2017; Incio *et al*., 2016; Ochi *et al*., 2012)^. Recent research found that PDAC cell-derived IL-1β is also play a critical role in promote PSC activation and tumor progression ^(Das *et al*., 2020b)^. Our data confirmed IL-1β can be expressed in PDAC cell lines and the secreted IL-1β in tumor conditional medium (CM) was significantly upregulated after treating with Gem, which may be the reason of secondary chemoresistance. Our data demonstrated that pancreatic cancer cell derived IL-1β can induce ESE3 overexpression in PSC by activating NF-κB signal pathway. NF-κB (p65) directly binds to ESE3 promoter and activates its expression. Silverman reported that IL-1β upregulated ESE-3 mRNA expression by activating MAPK pathway in bronchial smooth muscle cells ^(Silverman *et al*., 2002)^. Our results uncovered a new mechanism of IL-1β in inducing ESE3 expression in PSC.

In vitro experiment using cell lines demonstrated that conditional medium from ESE3 overexpressing PSC promoted PDAC cell survival, chemoresistance and migration. In vivo mouse model was confirmed the function of ESE3 (PSC) in promoting tumor growth and fibrosis. Furthermore, our data demonstrated that the mRNA and protein level of α-SMA, Collagen1 and IL-1β were significantly upregulated by ESE3. Mechanically, ESE3 directly binds to the promoter regions of α-SMA, Collagen1 and IL-1β and activates their expression. It is reported that α-SMA, Collagen1 and IL-1β overexpression are the prominent features of PSC activation ^(Sinn *et al*, 2014; Thomas & Radhakrishnan, 2019)^. PDACs and PSCs derived-IL-1β induce ESE3 overexpression to promote PSC activation and upregulation of α-SMA, Collagen1 and IL-1β. Importantly, ESE3 overexpression in PSC expression was an independent risk factor for both OS and DFS among PDAC patients.

Taken together, these data demonstrated a critical IL-1β/ ESE3(PSC)/ IL-1β positive feedback loop driving fibrosis, chemoresistance, and PDAC progression. Inhibiting this positive feedback loop might represent a novel strategy to reduce tumor fibrosis and to increase chemotherapeutic efficacy in PDAC.

## Materials and methods

### Immunohistochemistry (IHC)

With approval from the Ethics Committee, PDAC samples were obtained from 107 patients (aged 26 years to 78 years) undergoing surgical resection with histologic diagnosis of PDAC at the Tianjin Medical University Cancer Institute and Hospital. The patients’ histopathologic and clinical characteristics are shown in Table S2. Immunohistochemistry for ESE3 of PDAC patient tissues was performed using a DAB substrate kit (ZSGB-BIO, Beijing, China). Immunoreactivity was semi-quantitatively scored according to the estimated percentage of positive tumor cells as previously described. Staining intensity was scored 0 (negative), 1 (low), 2 (medium), and 3 (high). Staining extent was scored 0 (0% stained), 1 (1%-25% stained), 2 (26%-50% stained), and 3 (51%-100% stained). The final score was determined by multiplying the intensity scores with staining extent and ranged from 0 to 9. Final scores (intensity score × percentage score) less than 5 were considered as negative staining (−) or low staining (+), 5-9 were medium staining (++) or high staining (+++) ^(Zhao *et al*., 2017)^.

### Immunofluorescence (IF)

To assess ESE3 and α-SMA distribution, human PSCs were seeded onto glass slides for different treatments. The cells were then washed once with PBS and fixed with 4% paraformaldehyde in PBS for 15 min, permeabilized with 0.1% Triton X-100 in PBS for 30 min at room temperature, blocked for 1 h with 3% BSA in PBS. Then cells were stained with anti-ESE3 and α-SMA antibody (1:200 dilution, overnight at 4°C). Cells were mounted with DAPI Fluoromount-G media with DAPI nuclear stain (Southern Biotech). Slides were viewed with Olympus microscopy.

### PDAC cell culture and human and mouse primary PSC isolation and culture

Human PDAC cell lines, MIA-PaCa-2, BxPC-3, SW1990 and HEK 293 cell were obtained from the Committee of Type Culture Collection of Chinese Academy of Sciences (Shanghai, China). The human PDAC cell line L3.7 was a gift from Prof. Keping Xie (MD Anderson Cancer Center, Houston, TX). The murine PDAC cell line Panc02 was a gift from Shari Pilon-Thomas (Moffitt Cancer Center, FL). All the cell lines were recently authenticated in 2018 through the short tandem repeat analysis method. These cells were grown at 37°C in a humidified atmosphere of 95% air and 5% CO_2_ using Dulbecco’s modified Eagle medium (DMEM) with 10% fetal bovine serum (FBS).

The human primary PSCs (hp PSC) were isolated by the outgrowth method. PDAC surgical specimens obtained from patients at Tianjin Medical University Cancer Institute and Hospital were immediately cut into small pieces with a sharp scalpel. The PDAC tissue blocks were attached to the bottom of the 10cm dish, and carefully add 3ml of DMEM1/2 medium containing 20% FBS, 100 U/ml penicillin and 100 μg/ml streptomycin around the blocks. PSC were identified by immunofluorescence of α-SMA staining ^(Tian *et al*, 2019)^.

Mouse primary PSCs (mp PSC) were isolated from the pancreas of healthy C57BL/6 mice (3 months old) by collagenase digestion followed by Nycodenz® (Nycomed, Oslo, Norway) density gradient centrifugation ^(Jaster *et al*, 2002)^. Afterwards, the PSC were cultured in DMEM1/2 medium containing 20% FBS, 100 U/ml penicillin and 100 μg/ml streptomycin. The human PSCs were immortalized by transfection with SV40 large T antigen and human telomerase (hTERT) ^(Jesnowski *et al*, 2005)^.

### siRNA duplexes, plasmid constructs, transient transfection, stable transfection in pancreatic cancer cells

Small interfering RNAs (siRNAs) again ESE3, NF-κB were designed and synthesized from GenePharma (Shanghai, China) (Table S3). The human ESE3 cDNA was cloned into the pCDH plasmid expression vector.

ESE3 overexpression in PSC, Lentivirus-mediated plasmid was done using the pCDH-cDNA system (Biosettia) following the manufacturer’s instructions. Lentivirus encoding DNA were packaged as previously described ^(Elegheert *et al*, 2018)^. Following transfection, the medium containing lentivirus was collected, filtered, and transferred onto PSC. Infected cells were selected with puromycin (1 μg/mL) for 1 day.

For transfection, cells were plated at a density of 5×10^5^ cells/well in 6-well plates with serum-containing medium. When the cells were 80% confluent, the siRNA duplexes or overexpression plasmids were transfected into cells using Lipofectamine-2000 (Invitrogen) for 48 h. The cells were collected for cell migration analysis, Western blot analysis, and RT-PCR, apoptosis and cell cycle assay, etc.

### Western blot analysis

Whole-cell extracts were prepared by lysing cells with RIPA lysis buffer supplemented with a proteinase inhibitor cocktail (Sigma). The nuclear and cytoplasmic proteins of PSC were extracted according to the instructions of the nuclear-cytoplasm extraction reagents (Thermo Fisher Scientific, USA, Catalog number: 78835). Protein concentrations were quantified using Pierce protein assay kit (Pierce). Protein lysates (20 μg) were separated by SDS-PAGE, and target proteins were detected by Western blot analysis with antibodies (Table S3). Specific proteins were visualized using an enhanced chemiluminescence detection reagent (Pierce).

### Reverse-transcription polymerase chain reaction (RT-PCR)

Total RNA was isolated from transfected cells with TRIzol Reagent (Invitrogen) and used for first-strand cDNA synthesis using the First-Strand Synthesis System for RT-PCR (Takara). Each sample was processed in triplicate, and GAPDH was used as loading control. Each experiment was repeated independently for at least three times. PCR primers used are indicated in Table S3.

### PDAC-PSC co-culture, apoptosis and cell cycle assay, cell migration and wound healing assay

PDACs indirectly co-cultured with immortalized human PSC (ihPSC). Collect supernatants of pCDH-Vector ihPSC and pCDH-ESE3 ihPSC as the conditional medium (CM) and culture PDAC cell lines with the CM. 2×10^5^ the PDAC cells resuspended in DMEM medium containing 10% FBS were seeded in the lower chamber and 2×10^5^ pCDH-Vector ihPSC or 2×10^5^ pCDH-ESE3 ihPSC resuspended in 500μL of medium containing 10% FBS was placed in the upper chamber (Transwell chambers, Corning, USA). Then the MIA-PaCa-2 and BxPC-3 cell lines were treated with Gemcitabine (Gem). The apoptosis rate and cell cycle distribution were analyzed by flow cytometry. The Gem was obtained from Tianjin Medical University Cancer Institute and Hospital (Gemzar, Eli Lilly and Company).

The cell migration ability was measured using Transwell chambers (pore size, 8.0 μm, Corning, USA). 2 x10^5^ the PDAC cells resuspended in DMEM medium containing 2% FBS were seeded in the upper chamber and 2×10^5^ pCDH-Vector ihPSC or 2×10^5^ pCDH-ESE3 ihPSC resuspended in 500μL of medium containing 10% FBS was placed in the lower chamber. The cells were incubated for different times and the cells migrated to the bottom of the chamber were stained by three-step staining (Thermo Scientific). Cell migration was determined by counting the stained cells under a light microscope in 10 randomly selected fields. All experiments were repeated independently for at least three times. MIA-PaCa-2 and BxPC-3 cell lines were cultured with CM from pCDH-Vector ihPSC and pCDH-ESE3 ihPSC. Wound healing assay was performed according to published protocol.

### Animal studies in subcutaneous pancreatic cancer mouse model

Female 4-week-old nude nu/nu mice were maintained in a barrier facility on HEPA-filtered racks. All animal studies were conducted under an approved protocol in accordance with the principles and procedures outlined in the NIH Guide for the Care and Use of Laboratory Animals. Cells were harvested by trypsinization, washed in PBS, resuspended in a 1:1 solution of PBS/Matrigel, and 1×10^6^ L3.7+1×10^6^ pCDH-Vector ihPSC or pCDH-ESE3 ihPSC were injected subcutaneously into two flanks of 6 nude nu/nu mice. Primary tumors were measured in 3 dimensions (a, b, c), and volume was calculated as abc×0.52. Part of the tumor samples were fixated by formalin and embedded using paraffin and analyzed using H&E and IHC staining.

### Chromatin immunoprecipitation assay (ChIP) and Dual-luciferase assay

Chromatin immunoprecipitation assay was performed using a commercial kit (Merck-Millipore) according to the manufacturer’s instructions. The PCR primers are indicated in Table S3. Genomic DNA fragments of the human ESE3, α-SMA, Collagen1 and IL-1β genes, spanning from +1 to −2000 relative to the transcription initiation sites were generated by PCR and inserted into pGL3-Basic vectors. All constructs were sequenced to confirm their identity. Luciferase activity was measured using the Dual-Luciferase Reporter Assay System (Promega) according to the manufacturer’s instructions.

### Enzyme-linked immunosorbent assay (ELISA)

We collected conditional medium (CM) of Gem-treated PDAC cell lines and centrifuged at 1500 rpm for 5 min. The supernatants stored at −80°C until use. The production of IL-1β in supernatant were performed using ELISA kit according to the manufacturer’s instructions (Proteintech, China, Catalog number: KE00021).

### NF-κB p65 transcription factor assay

NF-κB p65 activity was determined by p65 binding ELISA (Abcam, ab133112). Binding ELISA kit was performed according to the manufacturer’s protocol using nuclear lysates and values were normalized to μg protein as determined by protein estimation (Abcam, ab113474).

### Statistical analysis

Two independent samples t-test was used for comparison between the two groups. The chi-square test was used to correlate the expression level of ESE3 in PSC with clinicopathological data. Kaplan-Meier method was used to estimate the overall survival (OS) or disease-free survival (DFS) of different ESE3 expression in PDAC. The log-rank method was used to compare the difference in OS/ DFS. The Cox proportional hazards regression model was used to assess the relationship between ESE3 expression levels in PSC and OS/ DFS. Each experiment was conducted independently for at least three times, and the data were presented as mean ± standard deviation (Mean ± SD). All tests presented were two-tailed and *P* < 0.05 was considered statistically significant. Statistical analysis was performed using SPSS 20.0 software and GraphPad Prism software.

## Authors’ contributions

TSZ, DX, FJJ, HWW and JHH designed, edited and led out the experiments of this study. TSZ, DX, FJJ, HWW, WRC, CBH, XCW, SG, JL and JHH conducted the experiments, data analysis, and critical discussions of the results. JHH provided material support and study supervision. SYY revised the manuscript. All authors contributed to the writing and editing of the manuscript and approved the final draft of the manuscript.

